# finsyncR, an R package to synchronize 27 years of fish and invertebrate data across the United States

**DOI:** 10.1101/2024.02.22.581615

**Authors:** Michael B. Mahon, Devin K. Jones, Ryan A. Hill, Terry N. Brown, Ethan A. Brown, Stefan Kunz, Samantha L. Rumschlag

**Affiliations:** Great Lakes Toxicology and Ecology Division, Center for Computational Toxicology and Exposure, U.S. Environmental Protection Agency, Duluth, MN, USA; Department of Biological Sciences and Environmental Change Initiative, University of Notre Dame, Notre Dame, IN, USA; Data Gathering and Analysis Division, Office of Pollution Prevention and Toxics, U.S. Environmental Protection Agency, Washington D.C., USA; Pacific Ecological Systems Division, Center for Public Health and Environmental Assessment, U.S. Environmental Protection Agency, Corvallis, OR, USA; Scientific Computing and Data Curation Division, Center for Computational Toxicology and Exposure, U.S. Environmental Protection Agency, Duluth, MN; Institute for Environmental Sciences, RPTU Kaiserslautern Landau, Landau, Germany

## Abstract

The United States government collects some of the most comprehensive data on freshwater systems in existence, but barriers exist for new users to apply these data without institutional guidance. ‘finsyncR’ (**f**ish and **in**vertebrate **sync**hronizer in **R**) is an R package that streamlines the process of acquiring, processing, and integrating fish and macroinvertebrate datasets collected in streams and rivers, increasing access to cleaned data and ensuring straightforward application of the data. The data sources are the US Environmental Protection Agency’s National River and Streams Assessment and US Geological Survey’s BioData. The resulting datasets span 27 years (1993 to 2019) and include 8,115 sites, 15,169 sampling events, 963 macroinvertebrate genera, and 687 fish species. We document common challenges to working with these data including understanding sampling designs, harmonizing taxonomy, calculating densities and standardized abundances, accounting for differences in sampling effort, and considering improvements in taxonomic identifications. We anticipate this package will spur research exploring changes in fish and macroinvertebrate communities across space and time and the causes and consequences of those changes.

## Background & Summary

United States (US) federal biomonitoring programs exist to assess the status and trends of freshwater ecosystems and communities^1^. These programs use standardized sampling protocols to produce data that are some of the most comprehensive in terms of spatial extent, temporal scope, and diversity of taxa surveyed. These data are used by the US government in assessment frameworks, but a wealth of additional knowledge can be gleaned from unique applications of these data. For instance, scientists have recently used data from federal biomonitoring programs to document long-term changes in stream biodiversity at a national scale^2^ and to test ecological theory on the influence of environmental change on species coexistence^3^. While most data associated with these biomonitoring efforts are publicly available, barriers remain for new, outside users to apply these data without institutional guidance. We have developed an R package, ‘finsyncR’ (**f**ish and **in**vertebrate **sync**hronizer in **R**), that flexibly acquires, processes, and integrates fish and macroinvertebrate biomonitoring datasets in rivers and streams of the US (Fig. 1). Here, we document the nuances and challenges of working with these data to dismantle barriers. The output datasets of the R package are sampling by taxa community matrices for fish and macroinvertebrates through time. Sites can also be linked to land use data within watersheds or catchments. The resulting output datasets span 27 years (1993 to 2019) and include 8,115 total unique sites (4,918 for fish and 7,789 for macroinvertebrates), 15,169 sampling events (8,193 for fish and 12,743 for macroinvertebrates), and 963 macroinvertebrate genera and 687 fish species (Fig. 2 and 3).

**Figure 1.**
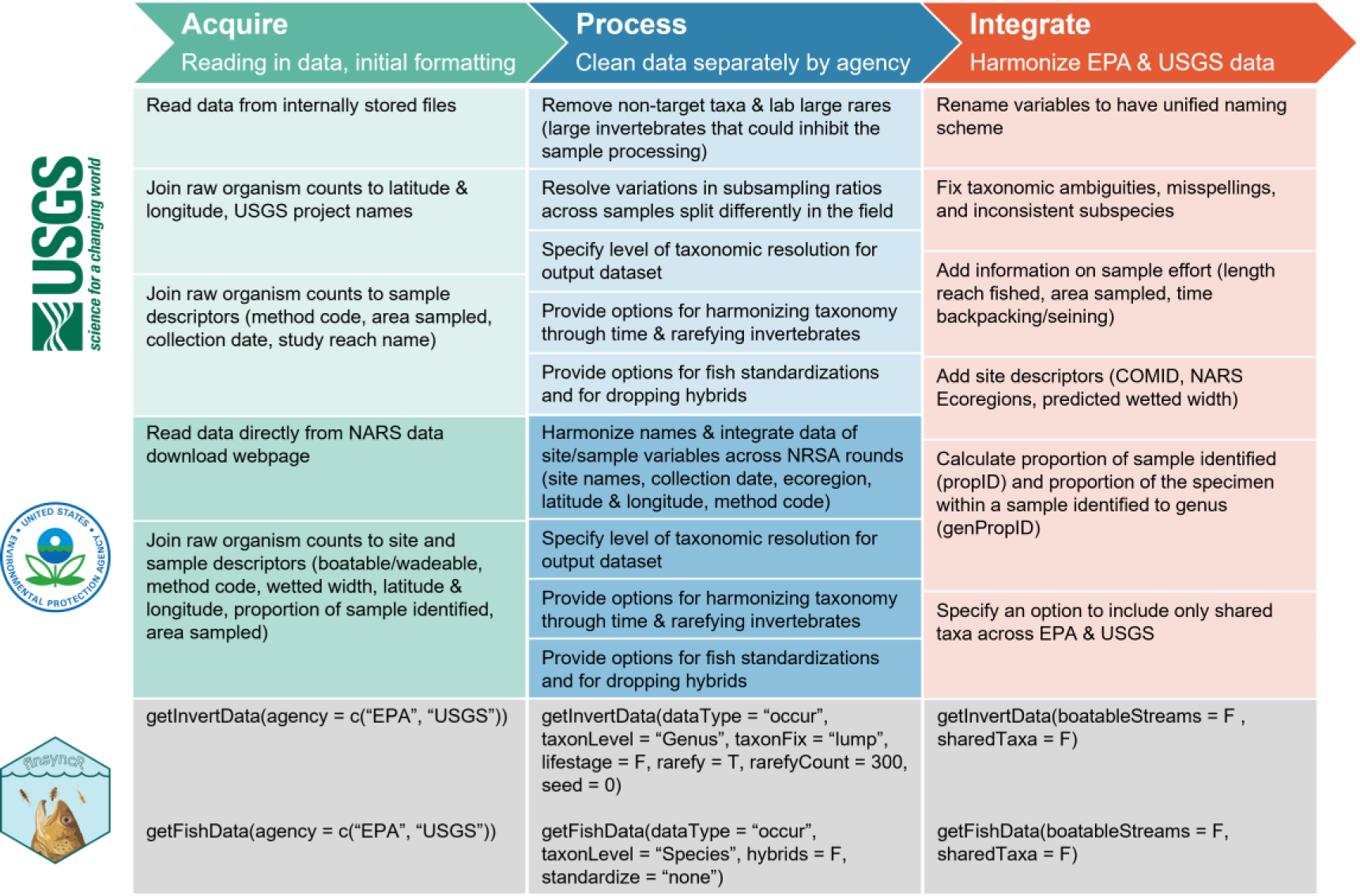
Workflow diagram depicting the construction of macroinvertebrate and fish functions. Function parameters are provided below each data management step in which they are developed. The metadata for the output datasets with definitions for variables included in this workflow can be accessed within the package using the code `vignettes("Metadata", "finsyncR")`.

**Figure 2.**
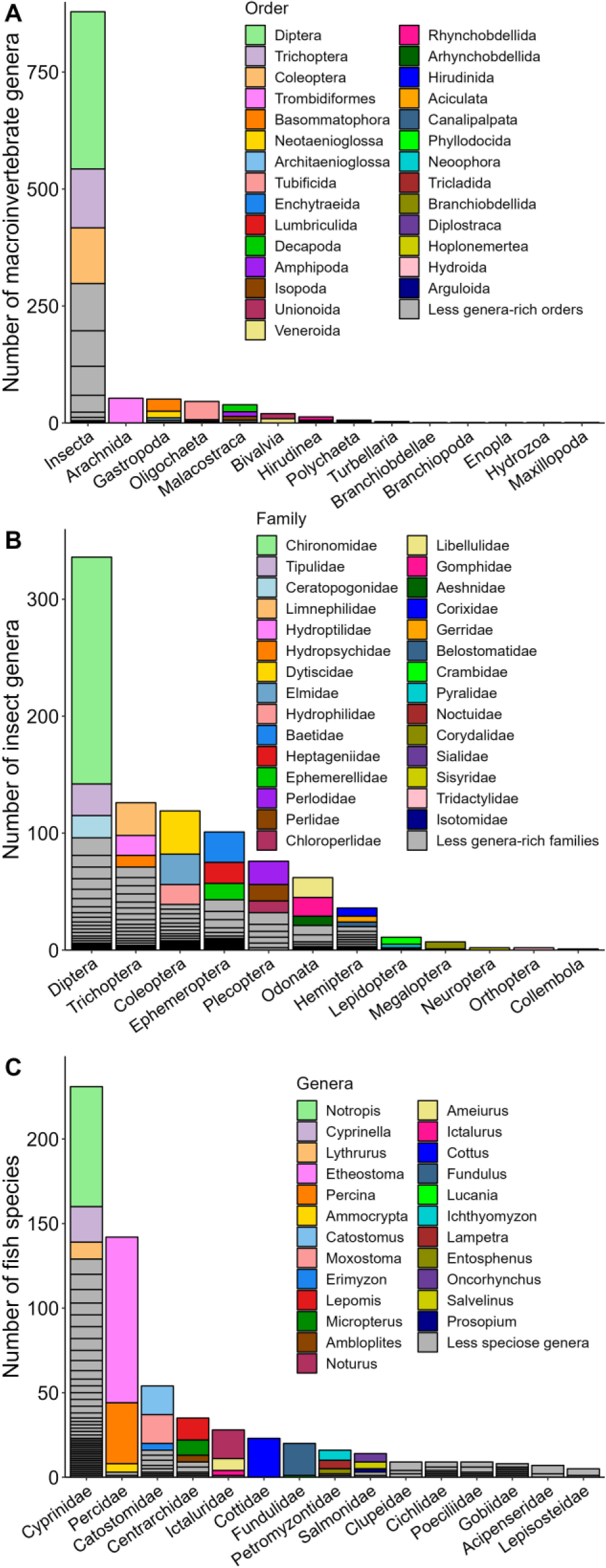
Plots showing the diversity of macroinvertebrate and fish. **A)** The number of macroinvertebrate genera within orders and phyla. **B)** The number of insect genera within families and orders. Given that the insecta phyla is diverse within macroinvertebrates, only insects are included to depict finer levels of diversity within this phylum. **C)** The number of fish species within genera and families. The first three most diverse **(A)** orders within phyla, **(B)** families within orders, and **(C)** genera within families are colored for ease of visualization.

**Figure 3.**
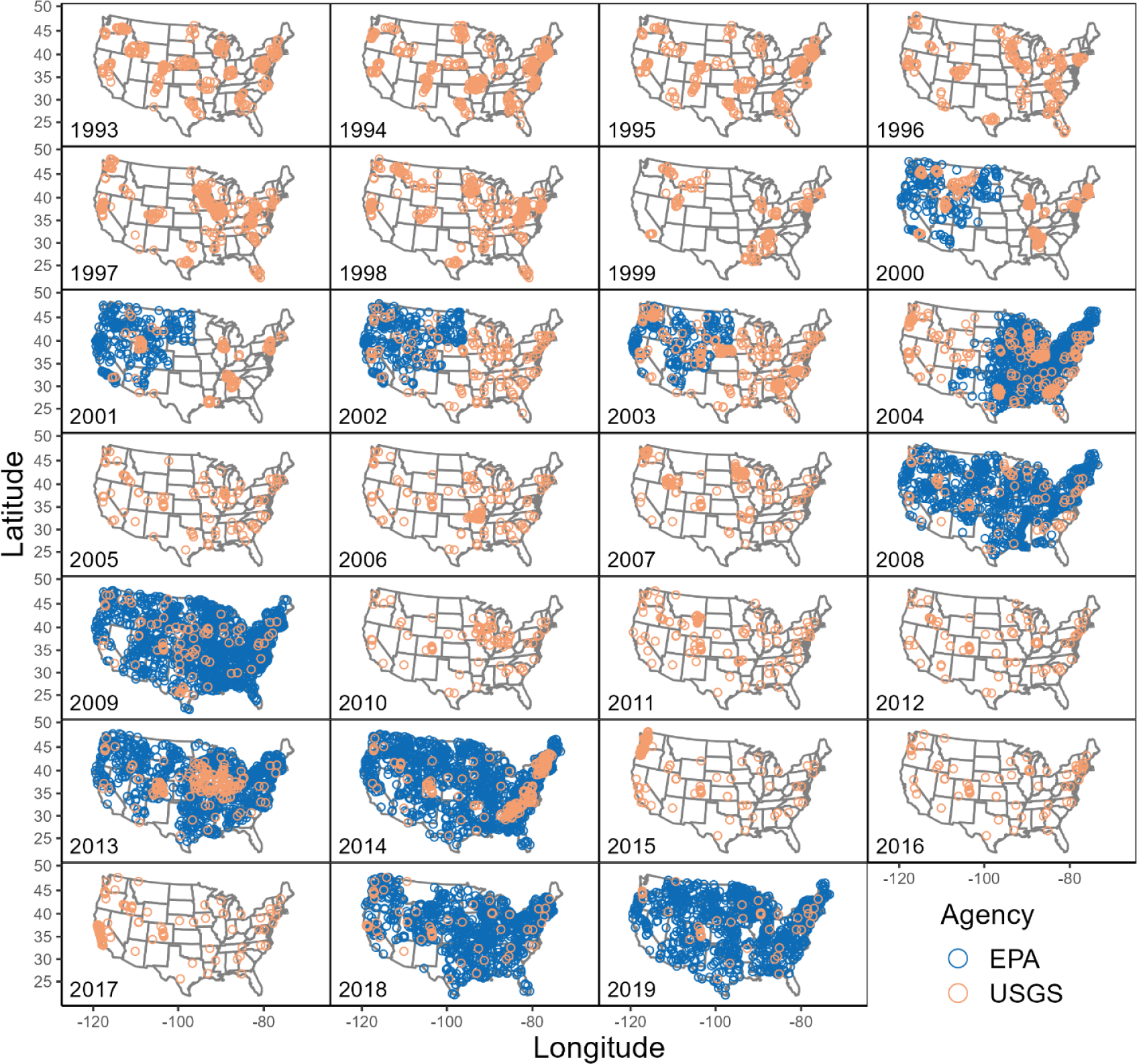
EPA and USGS sampling locations through years for both fish and macroinvertebrates. Sampling locations for USGS are set by research priorities of a given time period and assessment, and those priorities change through time. WSA, which is part of NRSA from 2000-2004, surveyed only wadeable streams in either eastern or western portions of the US in each year. In all other rounds of the NRSA, a randomized sampling strategy is used across the US, and surveys includes both wadeable and boatable waterbodies resulting in data that is representative of rivers and streams across the nation.

### Spatial Sampling Designs and Sampling Frequencies

The ‘finscyncR’ package integrates two federal biomonitoring datasets: the US Environmental Protection Agency’s (EPA) National Aquatic Resources Survey (NARS) and US Geological Survey’s (USGS) BioData. Operating under the directive of the Clean Water Act, NARS is a collaborative program between the EPA, states, and tribes tasked with assessing the state of the nation’s waters^1^. NARS surveys coastal waters, lakes, wetlands, and rivers and streams. Uniquely, NARS uses a spatially balanced randomized sampling design to ensure that data collected are representative of water bodies across the contiguous US^4^. In national assessments, each site is given a weight that is the number of miles the site represents relative to the contiguous US. Weights can be used to create population level estimates of measurements at a national scale. The surveys of rivers and streams is called the National Rivers and Streams Assessment (NRSA). Sampling rounds of NRSA occur every five years, and each round of sampling occurs across two years (Fig. 3). The exception is an early iteration of NRSA, called the Wadeable Streams Assessment (WSA), that surveyed only wadeable streams in either eastern or western portions of the US in each year from 2000 to 2004 (Fig. 3).

BioData is a database housing aquatic bioassessments and is run by the USGS. BioData includes sampling from national and regional assessments, including the National Water Quality Assessment, Regional Stream Quality Assessments, as well as state and local assessments. Sampling locations for USGS are set by research priorities of a given time period and assessment, and those priorities change through time. The BioData sampling design results in a clustered pattern in some early years, and in later years sites are dispersed throughout the US (Fig. 3). The strength of the combined datasets from the EPA and USGS is in their spatial coverage rather than the number of years an average site has been sampled. Most sites across both EPA and USGS are sampled for a single year although subsets of sites within each agency have been resampled across years (Fig. 4).

**Figure 4.**
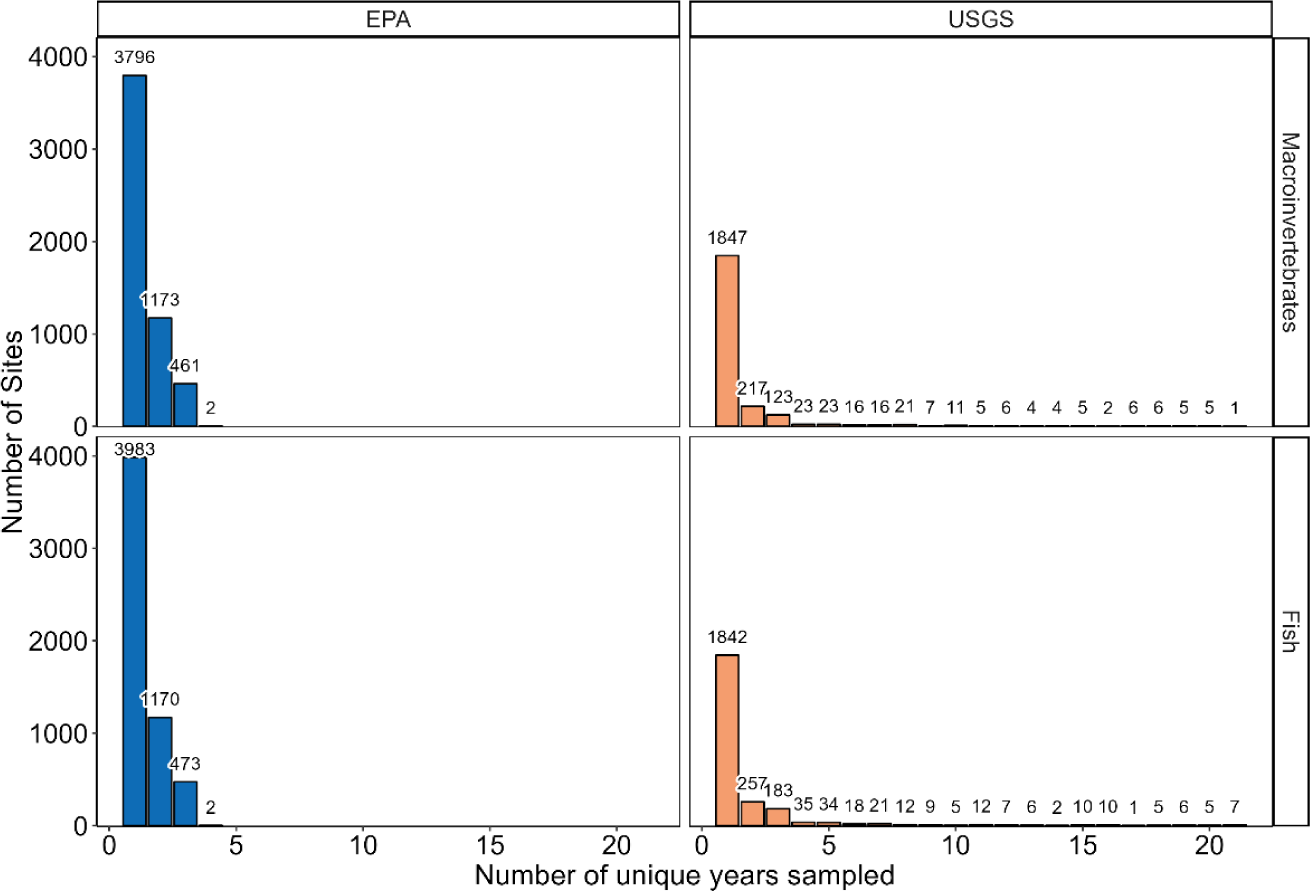
The distributions of the number of unique years sites were sampled for macroinvertebrates and fish by EPA and USGS. The vast majority of sites are sampled once. About 10% of EPA sites are resampled within a two-year sampling period (e.g. 2008/2009), and about 40-50% of sites are targeted for resampling across consecutive sampling periods (e.g. 2008/2009 to 2013/2014). However, the sites that are resampled between each set of consecutive rounds are not the same, so the 40-50% of 2008/2009 sites resampled in 2013/2014 are not the same 40-50% of 2008/2009 sites resampled in 2018/2019. There are 60 sites with macroinvertebrates sampled in 10 or more years and 76 sites with fish sampled in 10 or more years from the USGS.

The goals of the ‘finsyncR’ package are to: 1) unify EPA NRSA and USGS BioData creating a single national-scale, long-term dataset for fish and macroinvertebrates, 2) to streamline the process of acquiring, processing, and integrating these data, 3) to increase access to cleaned data and make the application of these data straightforward for users, and 4) to help users forgo common issues in data management by documenting the barriers to data use and by providing flexible strategies to overcome these barriers. While most data associated with these biomonitoring efforts are publicly available, barriers remain for new users to apply these data without institutional guidance. These barriers include understanding spatial sampling designs and sampling frequencies, processing and integrating datasets across years and federal agencies, accounting for differences in taxonomy between agencies and through time, calculating densities and standardized abundances from field counts of organisms, accounting for differences in sampling effort across sampling events, and accounting for improvements in taxonomic identifications through time. In this manuscript and associated package documentation and vignettes, we discuss these barriers and strategies to overcome them, so users can make informed decisions about the data given their specific research questions. With this information, new users will be able to conduct robust analyses and avoid making inferences based on known underlying data artifacts. We anticipate that this package and its associated documentation will ease the barriers to using some of the most comprehensive data on biotic communities within streams and rivers in existence, which could spur research advancements in streams and rivers. Potential future uses of the data include studies of changes in fish and macroinvertebrate communities across space and time and the environmental causes and ecological consequences of those changes.

## Methods

### Sampling Methods Summary

The EPA and USGS samples fish and macroinvertebrate communities during the summer months and low-flow conditions (generally, June to September). Macroinvertebrate sampling used a standardized targeted habitat approach of defined areas (typically 1.022 m^2^ for EPA wadeable sites, 3.353 m^2^ for EPA boatable sites, and 1.25 m^2^ for USGS sites) using a 500-µm mesh net^5–7^. The EPA macroinvertebrate samples were composite samples collected with a D-frame net along 11 equidistant transects within wadeable streams and from nearshore areas of transects at boatable streams^6^. The USGS macroinvertebrate samples were collected from five 0.25 m^2^ Slack samplers, typically from riffles, main-channel boundaries, and natural-bed features^5^. Only main-channel samples were included in the package because other samples include snags and woody debris, which likely influence the composition of organisms sampled. Additionally, sampling effort was more consistent across main-channel samples compared to other sample types. Identification of macroinvertebrates was typically limited to a target count. Target counts were 500 individuals for EPA samples and range from 100-600 for USGS samples. Macroinvertebrates were generally identified to genus in EPA samples and species in USGS samples. However, the package does not provide USGS species-level identifications for two reasons. First, we limit USGS identification to the genus-level to make the EPA and USGS data comparable. Second, species-level identifications of some groups of macroinvertebrates are inherently difficult to make^8^, so we limit identifications to the genus-level to avoid potentially inaccurate identifications.

Fish sampling used seining and/or electroshocking via backpack, barge tow, or boat. Nearly all of the EPA samples were collected by electroshocking. Most of the USGS samples were collected by electroshocking except for ∼20% by seining. The specific sampling method used was dependent upon the size of the waterbody and program and is denoted in the output datasets^5,7,9–13^. For EPA samples, in streams <12.5 m wide, reach length fished by electroshocking was a minimum of 150 m or 40 channel widths. At boatable sites and wadeable sites ≥12.5 m wide, the minimum reach length fished was 500 m or 20 channel widths. These sites were sampled until reach length fished equalled 40 channel widths or until 500 individual fish were collected. At some sites, no fish were collected despite attempts. These zero counts for fish are included in the output datasets from the package. For the USGS samples, reach length fished was a minimum of 150 m or 20 channel widths to a maximum of 300 m for wadeable streams. For USGS boatable streams, reach length fished was between 300 and 1000 m. Fish were collected using a two-pass protocol combining electrofishing and seine netting^5^. For all protocols organisms were enumerated and identified, and fish were released back into the stream alive. Fish that could not be identified in the field were retained for identification in the lab. The lowest level of taxonomic identification in the package is the species-level for fish because of inconsistent subspecies identifications. All sampling manuals associated with NRSA are available online (https://www.epa.gov/national-aquatic-resource-surveys/manuals-used-national-aquatic-resource-surveys).

### Acquiring Data

The code underlying the package follow three basic steps: acquiring, processing, and integrating the datasets (Fig. 1). Two functions, `getFishData()` and `getInvertData()`, generate sample by taxa matrices for fish and macroinvertebrates, respectively. The functions include the parameter `agency = c("USGS”, “EPA”)`that specifies which federal agencies the output datasets should be built from. The USGS data are called from locally hosted files (filepath: ∼/finsyncR/inst/extdata/), and EPA data are downloaded directly from the public facing EPA NARS website. Following data loading, USGS fish and macroinvertebrate count data are linked to USGS project, site, and sample descriptor datasets, and EPA fish and macroinvertebrate count data are linked to EPA site and sample descriptor datasets.

Next, some initial data processing occurs. Within the USGS macroinvertebrate samples, “lab large rare” organisms, which are large organisms, including crayfish, that inhibit uniform distribution of other organisms within a subsampling frame, are removed^5,7^. The package unifies EPA site names across sampling rounds using the NRSA “master site ID”, harmonizes variable names, and links sample data across sampling rounds.

### Processing Data across Years and Federal Agencies

The steps for data processing and integrating are based on dismantling barriers to data use. Data processing challenges are distinct between federal agencies and macroinvertebrate versus fish samples. For macroinvertebrates, barriers include reconciling multiple subsampling ratios within single samples, resolving inconsistent taxonomic resolution across organisms, and making organisms identified with a slash name (e.g. genus1/genus2) explicit in EPA samples.

First, a subset of USGS samples contain multiple values for the proportion of the sample identified, which is the result of splitting samples multiple times in the field or lab (for a description of splitting samples, see the ‘Calculating Densities and Standardized Abundances from Field Counts’ below). Eight samples have more than one field split ratio and are removed because multiple field split ratios could not be reconciled. For samples with multiple lab subsampling ratios, the package sums unique organism counts across the multiple ratios and takes a proportional mean of lab subsampling ratios based on organism counts.

Next, within the EPA dataset, there are instances of inconsistent taxonomic resolution, where some organisms are identified to subfamily or tribe, which is a level of taxonomic resolution that is not available across all macroinvertebrates. In these instances, the package reassigns the identification to the family level. Lastly, for EPA organisms identified with a slash genus (e.g. genus1/genus2), which are identifications given to an organism when multiple genera are indistinguishable, the package pulls the identification from the “target taxon” column instead of the “genus” column. The “genus” column is used for all other organisms identified with a single genus.

Within the fish dataset, barriers to dataset unification include differences in hybrid naming schemes, inclusion of non-target taxa, use of common names, and designation of stream size classes. The naming scheme for hybrid individuals differs between the USGS and EPA fish datasets. The naming scheme for USGS hybrids is “Genus species1 x Genus species2” and for EPA hybrids is “Genus1 x Genus2 species1 x species2”. The package ensures that all hybrids follow the EPA scheme, so that if parental genera are shared, the naming scheme is “Genus species1 x species2”. The package provides users the option to retain or remove all hybrid individuals from the generated dataset with the parameter `hybrid = FALSE`. Within the EPA dataset, non-target taxa, specifically amphibians, are included in some raw samples. The package removes non-target taxa. Additionally, only common fish names are provided in the EPA datasets. The package updates the common names to scientific names and provides the associated taxonomic information.

Finally, stream size classes (small wadeable, large wadeable, and boatable) are unified for USGS and EPA datasets based on sampling protocols and measured widths of streams. For USGS samples, the package classifies samples in which boating, snorkelling, gillnets, or beach seines were employed as boatable streams. The package classifies samples collected by barge/boat tow electroshocking as large wadeable streams. The package classifies samples collected by seine netting alone or backpack electroshocking as small wadeable streams^7^. For EPA samples, wadeable and boatable designations are provided for most sites. The package reclassifies EPA-designated wadeable streams as large or small, depending on the mean wetted width at the time of sampling. Streams with wetted widths <12.5m are small wadeable, and streams with wetted widths >=12.5m are large wadeable. In a small number of instances in which no wadeable or boatable designation is provided, the package classifies streams as small wadeable if the wetted width was <12.5m, as large wadeable if the wetted with was >=12.5m and <=25m, and as boatable if the wetted width was > 25m^12,13^.

### Accounting for Differences in Taxonomy between Agencies and through Time

Another major barrier to data use is taxonomic harmonization both between agencies and through time. The package automatically harmonizes macroinvertebrate taxonomy between the EPA and USGS and provides options for harmonizing taxonomy through time. Taxonomy is automatically harmonized in three ways to join USGS and EPA datasets. First, in instances when a given organism has been identified at the species level and the genus that was originally given has been updated to a new genus, the package updates the genus designation by cross walking the bench identification to the current genus. Second, there are instances when a slash genus was not used consistently between USGS and EPA. Slash genera are designations of multiple genera given to an organism during bench identification when multiple genera are indistinguishable (such as *Cricotopus/Orthocladius*). As a hypothetical example, the USGS data contains an organism identified as genus1/genus2, while EPA data included separate identifications of genus1 and genus2. To ensure consistency between the agencies, the package renames all single instances of genus1 and genus2 to genus1/genus2 when combining the EPA and USGS samples into a single dataset. Third, within the EPA dataset, a group of chironomids were identified to a genus-group (*Thienemannimyia* genus group), which consists of the chironomid genera *Arctopelopia*, *Conchapelopia*, *Hayesomyia*, *Helopelopia*, *Meropelopia*, *Rheopelopia*, *Telopelopia*, and *Thienemannimyia*. The package replaces instances of organisms identified to one of these individual genera or the *Thienemannimyia* genus group with a single slash genus, *Arctopelopia*/*Conchapelopia*/*Hayesomyia*/*Helopelopia*/*Meropelopia*/*Rheopelopia*/*Telopelopi a*/*Thienemannimyia*.

Across the time span of the USGS and EPA macroinvertebrate datasets, the taxonomy of many macroinvertebrates changed, specifically membership of species within genera. The package provides options to harmonize taxonomy through time, especially in instances in which membership of a given organism within a genus is not clear. In certain scenarios a given organism may have been identified to genus at the lab bench, yet when taxonomic updates occurred, only some species within the originally designated genus moved to a different genus. This scenario prevents identification of a given organism’s “true” genus because it is possible the organism in question might belong to the original genus or the new, updated genus. For instance, consider an organism identified with the hypothetical genus *Hereiam*. At the time of the identification, there were only two species within this genus: *Hereiam here* and *Hereiam there*. In the time since the identification at the lab bench, the taxonomy has been reorganized, and presently the genus *Hereiam* contains one species *Hereiam here*, and a new genus has been created. *Hereiam there* is now *Thereiam there*. So, it is unclear whether the organism, originally identified as *Hereiam* belongs to *Hereiam* or *Thereiam*. This issue is especially difficult to overcome when attempting to harmonize samples that were identified before and after a taxonomic update occurred.

The package provides three options for accounting for these changes in taxonomy through time using the parameter `taxonFix`. First, the default option, `taxonFix = “lump”` creates complexes of genera, by combining all possible genera for a given organism, hereafter termed “lumped genera”. This approach prioritizes retaining observations of organisms by giving a unified name (e.g. genus1/genus2/genus3) to a complex of genera that have been linked through changes in taxonomy through time. With this approach, there is no temporal increase or decrease in the number of genera, given that individual genera (e.g. genus1) never appear outside of the lump genera name (e.g. genus1/genus2/genus3). So, in the hypothetical example above, *Hereiam* and *Thereiam* would occur as *Hereiam/Thereiam* throughout the generated dataset. In total, 82 macroinvertebrate genera are linked by changes in taxonomy through time (Fig. 5). The “lumping” approach described above reduces these 82 genera to 11 lumped genera. All but three lumped genera were composed of two individual genera. A single lumped genera group within the order Ephemeroptera includes 54 genera, and two additional lumped genera of Ephemeroptera includes six genera each. Because of the complexity of Ephemeroptera taxonomy, careful consideration should be given to inferences that can be made when evaluating trends in Ephemeroptera diversity and abundance.

**Figure 5.**
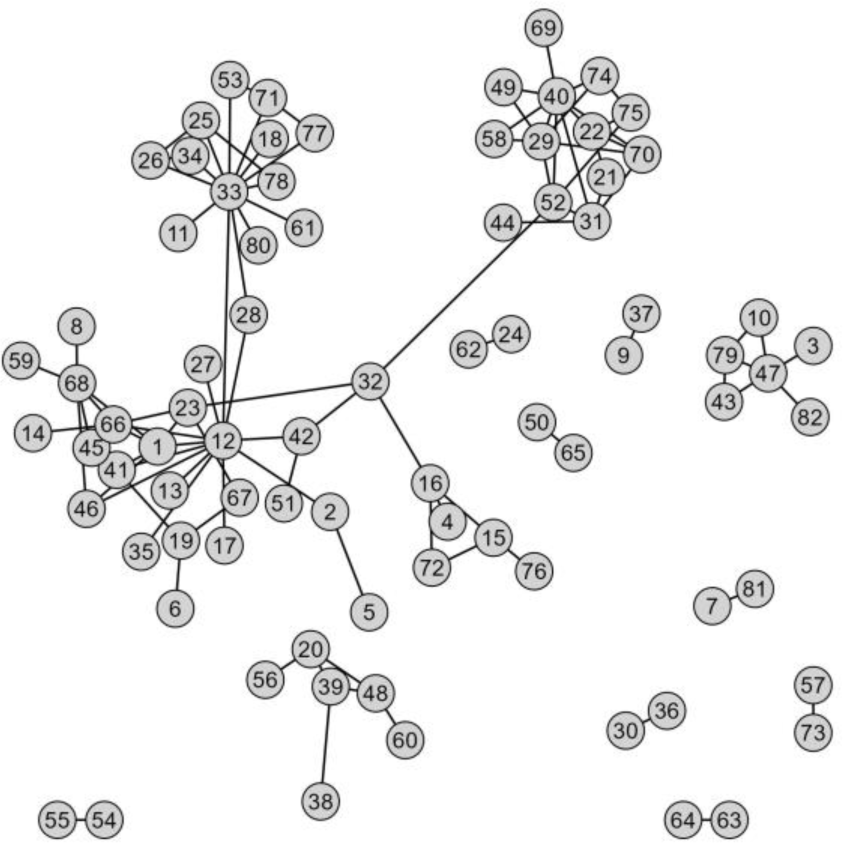
Relationships among genera linked by taxonomic reorganization through time. Each numbered circle represents an individual genus. Lines between genera indicate that the taxonomic designation of at least one species from one of the genus pairs was updated to be in the other genus in the pair. Lumped genera are formed by sets of these linked genera. For identities of these groups, how common each genera is in the USGS and EPA data, and their lowest common taxonomic designation see Table S1.

If `taxonLevel = “Mixed”`, which returns a macroinvertebrate dataset with the lowest level of taxonomic identification for all specimen, a user may wish to roll up the 11 lumped genera up to the lowest level of consistent taxonomic designation for each lumped genera group (Table S1). The function `lumpRollUp()` takes an input dataset produced by `getInvertData()` and performs this taxonomic transformation.

The second option, `taxonFix = “remove”`, prioritizes accurate identification by dropping observations that cannot be confidently identified to a single genus, which includes the lumped genera previously described. In addition, this option removes organisms identified using slash genera (e.g., genus1/genus2). So, the “remove” option results in a dataset that has fewer genera compared to dataset generated with the “lump” option. The "remove" option is the best option if accurate identification is a high priority for users.

Finally, `taxonFix = “none”` returns identifications in their original forms. The "none" option generates slash genera for organisms that are given a slash genus either at the time of identification or to harmonize occurrences of slash genera between USGS and EPA observations as described above. Care should be taken to harmonize taxonomy either with the approaches described above or by some alternative when generating long time scale datasets on the entire community of macroinvertebrates because changes in taxonomy can make it artificially appear as though some genera are either appearing or disappearing in time. The macroinvertebrate taxonomy dataset used to harmonize was provided by USGS BioData.

For fish, taxonomic harmonization includes fixing misspellings, removing inconsistent subspecies, and clarifying taxonomic ambiguities. Using the EPA fish taxonomy dataset, the taxize R package^14^, and Fishbase^15^, the package harmonizes the spellings of scientific names and updates taxonomies within the USGS and EPA datasets. The EPA includes subspecies that the USGS does not. So, to ensure compatibility across agencies, the package assigns species designation to organisms identified to the subspecies level. In both the USGS and EPA fish observations, there are instances of taxonomic ambiguities. When organisms are given “sp.” as the species name, suggesting difficulty in discerning the species, the package gives the organism the genus level identification. Additionally, when the scientific name given include “cf.” (*conferret*), which suggests uncertainty in the species identification, the organism is given the assumed species identification.

### Calculating Densities and Standardized Abundances from Field Counts

No density or standardized abundance measurements are provided in the original datasets which can be a barrier to data use. The package automates the calculation of macroinvertebrate densities and fish standardized abundances. For macroinvertebrates, density calculations are simply macroinvertebrate abundance divided by the area sampled (Fig. 6). Macroinvertebrate abundance is the number of individuals counted in a gridded tray in the lab multiplied by the inverse of the proportion of the sample collected in the field that was identified at the lab bench (*PropID*). The calculation of *PropID* differs by agency because of differences in splitting samples in the field. For USGS samples, a proportion of the homogenized field sample is separated before specimen identification and enumeration in the lab. This proportion is the “field split ratio” (Fig. 6). EPA field samples are not split. For both EPA and USGS the processing of samples in the lab is similar. In the lab, organisms in a sample are placed uniformly among the grids in a subsampling tray. Grids are randomly selected and organisms within a given grid are identified until a target organism count is reached. The proportion of grids used to identify macroinvertebrates at the lab bench is the “lab subsampling ratio”. For USGS samples, *PropID* is calculated by multiplying the “field split ratio” and the “lab subsampling ratio” (Fig. 6). For EPA samples, *PropID* is equal to the “lab subsampling ratio” (Fig. 6). The package does not provide abundance and raw counts of macroinvertebrates, but they can be back calculated from density. We provide code to do these back calculation in a vignette that can be accessed with the following code `vignettes("BackCalculation", "finsyncR")`. This vignette also includes code to rarefy raw counts of samples.

**Figure 6.**
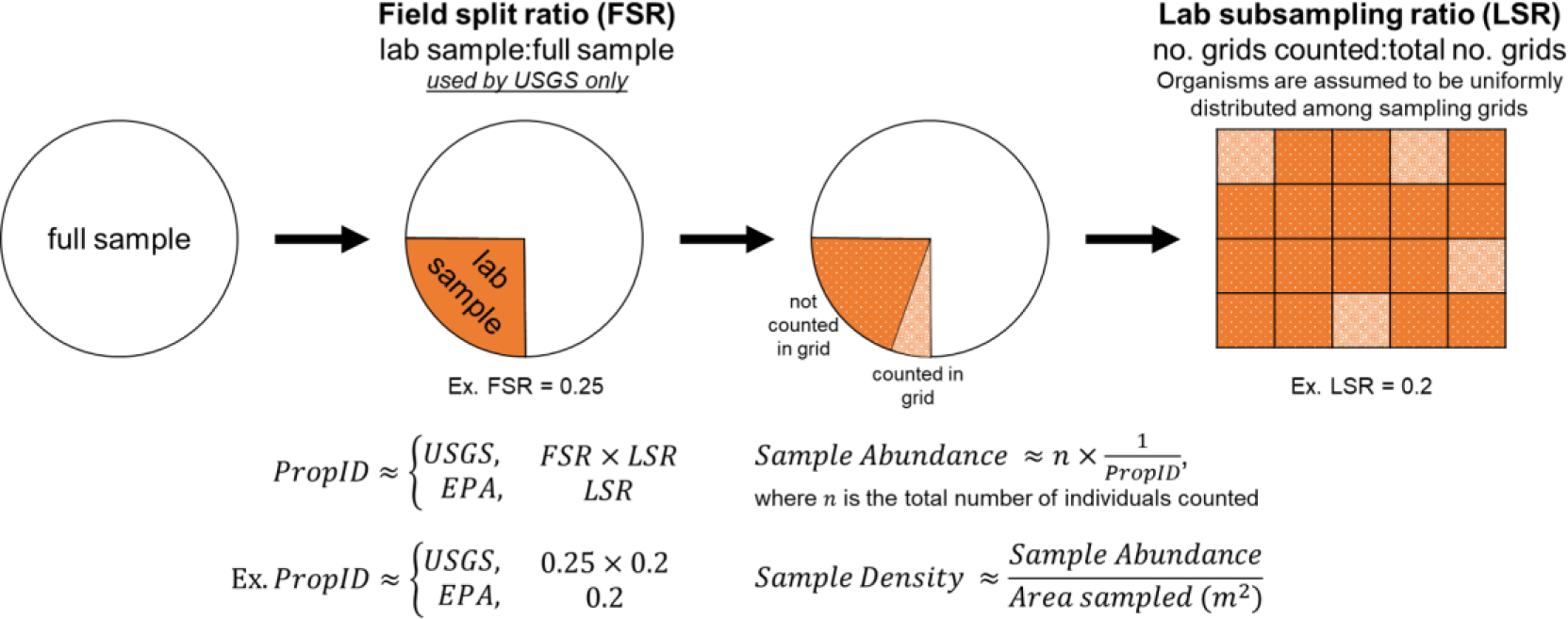
Calculation of sample densities for USGS and EPA samples. The field split ratio is the proportion of the field collected sample that is brought into the lab to be identified. Note that only the USGS data included field split ratios. Lab subsampling ratio is the proportion of grids in a subsampling tray used to identify and enumerate macroinvertebrates in the lab.

Within the fish datasets, sampling effort (e.g. minutes shocking, seine nets hauls, snorkelling transects, length of stream reach fished) and method (electroshocking, seining, and snorkelling) vary across samples and influence the number of fish per sample. The package provides users with the ability to account for differences in sampling effort and reach length fished in the calculation of standardized abundances. First, the package can calculate catch per unit effort (CPUE), which results in abundance values standardized by sampling effort and reach length fished:

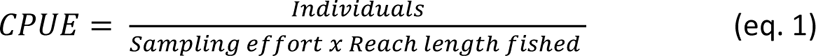

Second, to account for differences in efficacy among shocking, seine netting, and snorkelling, multigear mean standardization (MGMS) is another standardization method provided. For MGMS, individual species abundances are standardized for each gear type (e.g. electroshock, seine net, snorkel), as above in CPUE. Then, for each gear type, the CPUE of all *i* species in each sample *j* is summed, to get Total Catch Per Unit Effort for sample *j* (TCPUE*_j_*):

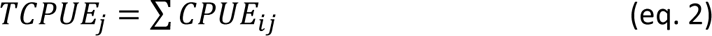

For each gear type, the mean TCPUE is calculated, 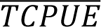. Next, to standardize each gear, CPUE for each species *i* is divided by 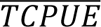:

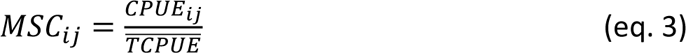

where MSC*_ij_* is mean standardized catch of species *i* in observation *j*. The units of sampling effort cancel in the calculation of MSC*_ij_*, and patterns of relative abundance of species within and across observations are preserved, which can be important for community analyses (e.g. relative evenness, Bray-Curtis distance calculation). The function then sums the MSC*_ij_* among gear types, resulting in a single row of data for each sampling event, regardless of the number of gear types used. See Gibson-Reinemer et al. (2017)^16^ for more information regarding the computation of MGMS.

### Integrating Data across Years and Federal Agencies

The package ensures consistency in naming and formatting site and sample variables across EPA and USGS fish and macroinvertebrate datasets. These site and sampling variables include agency, agency-specific site name, geographic coordinates, and collection date. The datasets also include information on sampling effort (macroinvertebrates: area sampled, proportion of sample identified, proportion of specimen identified to genus; fish: reach length sampled and sampling method effort).

The package also provides site-specific COMIDs from US National Hydrography Dataset Plus V2^17^ to allow users to link additional hydrologic data. In addition, the package provides Strahler stream order^18^ and wetted width as measures of stream size. Measured wetted width is only available for EPA data, so the package also provides predicted wetted width data gathered from StreamCat^19^ originally from Doyle et al. (2023)^20^. Finally, because of known ecoregional differences in species pools and sampling designs, the package provides NARS aggregated Omernick level III ecoregions for each site^21^.

### Linking Sampling Sites to Land Cover Data with getNLCDdata()

The package also contains a function, `getNLCDData()`, that retrieves land cover data from the National Land Cover Database (NLCD) for sampling sites at the catchment and watershed scales. The unit of land cover returned is percent area of catchment or watershed. This function directly accesses the EPA StreamCat API^19^ and gathers NLCD data based on sampling location from NHDPlus COMIDs and collection year. Users provide site names and collection year from the macroinvertebrate or fish datasets as input parameters to `getNLCDData()`. Then, the function matches sampling sites to NLCD data in time. NLCD data are available in the following years: 2001, 2004, 2006, 2008, 2011, 2013, 2016, and 2019. For instances of samples that do not have a match in collection year to a year of NLCD data, samples are matched to the closest year of NLCD data. For instance, a site sampled in 1995 is matched to 2001 NLCD data. For samples (e.g. collection year 2005) that fall at the midpoint of two NLCD years (e.g. 2004 and 2006 NLCD data), the function pulls NLCD data from the earlier year (e.g. 2004 NLCD data). Users can specify whether NLCD data should be extracted at the catchment or watershed scale. Catchments are the portion of the landscape where surface flow drains into an COMID stream segment, excluding any upstream contributions. Watersheds are the set of hydrologically connected catchments, consisting of all upstream catchments that contribute flow into any catchment^19^. Additionally, the package provides users with the ability to group NLCD data into five broad categories. These categories include “water” (all wetland and open water land covers), “urban” (all urban land uses), “forest” (deciduous, coniferous, and mixed forests land covers), “open” (shrub, barren land, grassland, and hay/pasture land covers), and “crop” (cultivated crop land use).

## Data Records

### Storage

We store the ‘finsyncR’ package via two different storage systems to provide users with alternatives to access and use the data. The FigShare repository (DOI: 10.6084/m9.figshare.24938607) stores the fixed, original version of the package (version 1.0.0). The EPA GitHub repository (github.com/USEPA/finsyncR) stores the future updated versions of the R package. Future updates may occur as new datasets are integrated, such as new cycles of NRSA sampling.

### Input Datasets

The raw sources of fish and macroinvertebrate data are currently the EPA NRSA and USGS BioData. Below we cite the original sources of fish and macroinvertebrate data. BioData are stored internally in the package (∼/Rlibrary/finsyncR/inst/extdata/). Most data from NRSA are directly downloaded by the package from the public website included in the data citations. NRSA data on sampling effort (for macroinvertebrates: proportion sample identified [*PropID*] and area sampled [*AreaSampled_m*], for fish: reach length fished [*ReachLengthFished_m*], sampling method [*SampleMethod*], sampling effort [*MethodEffort*]) are stored locally within the R package (∼/Rlibrary/finsyncR/inst/extdata/).

- U.S. Geological Survey, 2020. BioData - Aquatic bioassessment data for the Nation: U.S. Geological Survey database, accessed 17 December 2020, at https://doi.org/10.5066/F77W698B^22,23^
- U.S. Environmental Protection Agency. 2006. National Aquatic Resource Surveys. Wadeable Streams Assessment 2004 (data and metadata files). Available from U.S. EPA web page: https://www.epa.gov/national-aquatic-resource-surveys/data-national-aquatic-resource-surveys.^24^
- U.S. Environmental Protection Agency. 2016. National Aquatic Resource Surveys. National Rivers and Streams Assessment 2008-2009 (data and metadata files). Available from U.S. EPA web page: https://www.epa.gov/national-aquatic-resource-surveys/data-national-aquatic-resource-surveys.^25^
- U.S. Environmental Protection Agency. 2020. National Aquatic Resource Surveys. National Rivers and Streams Assessment 2013-2014 (data and metadata files). Available from U.S. EPA web page: https://www.epa.gov/national-aquatic-resource-surveys/data-national-aquatic-resource-surveys.^26^
- U.S. Environmental Protection Agency. 2022. National Aquatic Resource Surveys. National Rivers and Streams Assessment 2018-2019 (data and metadata files). Available from U.S. EPA web page: https://www.epa.gov/national-aquatic-resource-surveys/data-national-aquatic-resource-surveys.^27^

Land cover data from the National Land Cover Database (NLCD) is available for sampling sites at the catchment and watershed scales from the StreamCat dataset^19^. The function `getNLCDData()` directly accesses the StreamCat API to gather NLCD data based on sampling year and sampling location. More information on accessing StreamCat data through the API can be found here: https://www.epa.gov/national-aquatic-resource-surveys/streamcat-metrics-rest-api.

### Output Datasets

The function getFishData() produces standardized abundance (catch per-unit effort) and occurrence data for fish at various levels of taxonomic resolution. The function getInvertData() produces density and occurrence data for macroinvertebrates at various levels of taxonomic resolution. In both functions, the parameter `dataType` specifies whether the output is abundance-based or occurrence, and `taxonLevel` specifies the taxonomic resolution. Observations taxonomically coarser than the taxonomic level specified by the user are dropped from the output community matrix. For instance, if a user specified a taxonomic resolution of “genus”, then observations made at the subfamily or coarser level would be removed from the output dataset. Additionally, the getInvertData() function provides an option of returning the lowest level of taxonomic identification for all specimen by setting `taxonLevel = “Mixed”`.

The datasets can be flexibly produced using parameters of the functions to best suit the needs of the specific research question of the user. Some of the options provided by the parameters are described in this manuscript. The parameters are described in additional detail in the Getting Started Vignette, which can be accessed with the following code, `vignettes("Getting Started", "finsyncR")`.

Finally, the function getNLCDData() retrieves land cover data for fish and macroinvertebrate sampling sites at the catchment and watershed scales.

### Metadata for Output Datasets

The metadata for the output datasets can be accessed within the package using the code, `vignettes("Metadata", "finsyncR")`.

## Technical Validation

The technical validation of the ‘finsyncR’ package and the resulting datasets is based on (1) individual review procedures of input datasets, (2) comparability of the sampling methods across biomonitoring efforts, and (3) documentation of the sources of variation that should be considered in the context of addressing specific research questions.

### Individual Review Process of Input Datasets

USGS and EPA produced datasets are held to a high standard of quality. For instance, the input datasets included in ‘finsyncR’ package have been collected under federal quality assurance project plans. Additionally, before release to the public, federal data products are cleared through internal review procedures. As a result of these processes, the input data have already individually received technical validation.

### Comparability of Sampling Methods

The methods used by EPA and USGS are designed to characterize the entire fish and macroinvertebrate communities, suggesting they are comparable in theory. References to manuals describing the different methods used to collect fish and macroinvertebrates (sample type codes [*SampleTypeCode*]) are present in the metadata. The metadata for the output datasets can be accessed within the package using the code, `vignettes("Metadata", "finsyncR")`.

To ideally test method comparability, communities sampled at the same time and location using various methods should be compared. Unfortunately, these comparisons are not possible because sites sampled using this design do not exist in the present datasets. However, previous studies have compared some of the collection methods using this approach. For instance, some of the macroinvertebrate communities in wadeable streams were sampled with the following codes: BERW, IRTH, SWAMP, EMAP, CDPHE, and PNAMP. All methods use kick net samplers with similar mesh sizes, and studies suggest that methods are similar in their ability to detect the presence and absence of taxa^28–30^. Fish are collected through seining, snorkelling, backpack or raft electrofishing, or some combination. Electroshocking is the most common method across the dataset. Sampling methods used by EPA and USGS using electroshocking produce similar biotic indices for fish, suggesting methods are comparable^31^. Additionally, BioData includes USGS fish samples that were collected with NRSA methods, suggesting federal programs have confidence that methods are comparable enough for data to be combined^23^.

In addition, we directly compared the ability of different methods to detect occurrences of fish and macroinvertebrate taxa in output datasets. We conducted partial distance-based redundancy analyses (dbRDAs) of Bray-Curtis dissimilarities based on the presence and absence of macroinvertebrate genera and fish species at sites. We examined the amount of variance attributable to sampling methods (sample type codes) after accounting for spatial (HUC8) and temporal (collection year) variation. For the dbRDAs, fish samples collected by snorkelling were removed because only five samples had been collected by this method. Sampling method accounts for 2% of the total variation in macroinvertebrate communities and 1% of variation in fish communities (Fig. 7). For macroinvertebrates, ellipses of all sampling methods show high levels of overlap, suggesting different sampling methods collect comparable communities (Fig. 7A). For fish, EPA sampling methods are highly consistent regardless of stream size and fishing gear, demonstrating comparability of methods even across stream sizes (blue ellipses, Fig.7B). Across agencies electroshocking approaches overlapped, suggesting comparability of electroshocking methods. USGS sampling methods showed some small separations by seining and electroshocking samples (orange ellipses, Fig. 7B), but regardless of the degree of separation, sampling methods accounted for 1% of the total variation, which lends support to the suitability of the unification of samples across methods and agencies.

**Figure 7.**
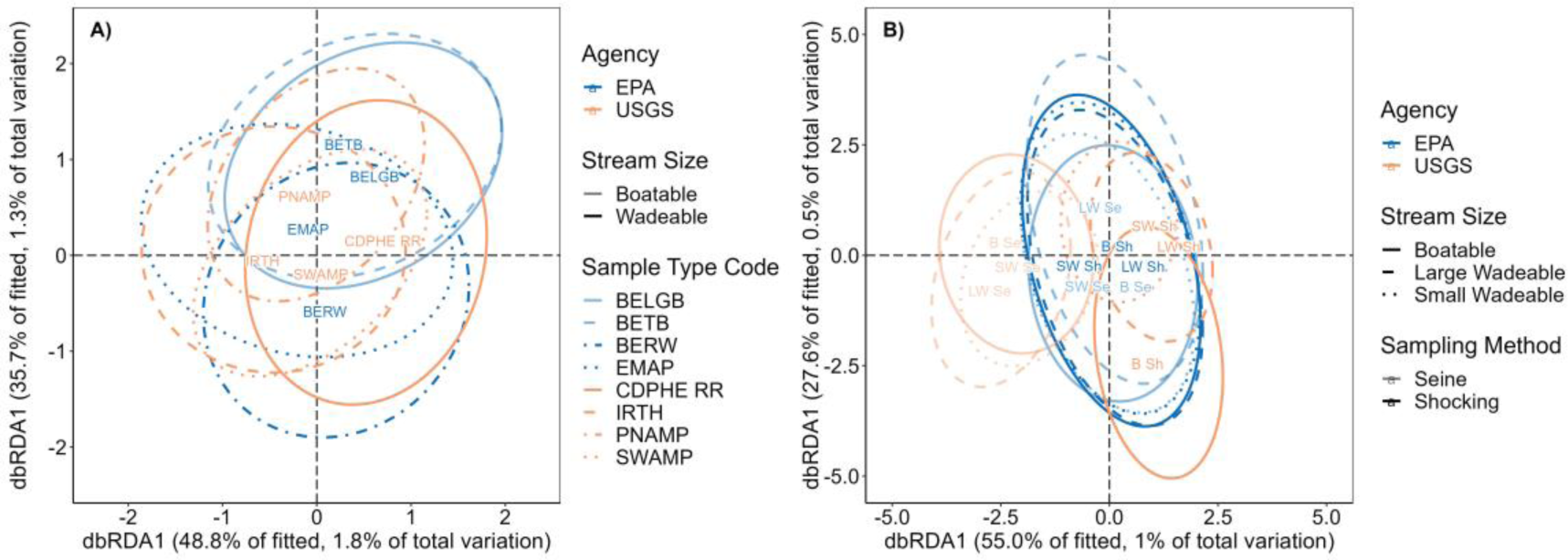
Comparability of sampling methods and stream size for sampling macroinvertebrate and fish communities. Partial distance-based redundancy analysis (partial dbRDA) of macroinvertebrate **(A)** and fish **(B)** community by sampling method, accounting for differences in HUC8 and collection year. Sampling method accounts for a small proportion of the total variation in the macroinvertebrate (2%) and fish (1%) datasets, and ellipses overlap across sampling methods, suggesting that the different sampling methods between and within agencies collect comparable communities, supporting the unification of these data. Each ellipse is one standard deviation from points within groups. **(A)** BERW combines sample type codes of “BERW”, “BERWW”, and “REACHWIDE”, which are NRSA sampling round-specific names for sampling methods in wadeable streams. Abbreviation definitions in **(A)** are USGS Invertebrate Targeted Habitat (IRTH), EPA Benthos Reach-wide (BERW), EPA Benthos Low Gradient Boatable (BELGB), EPA Benthos Transect Boatable (BETB), EPA Environmental Monitoring and Assessment Program (EMAP), Pacific Northwest Aquatic Monitoring Partnership (PNAMP), Colorado Department of Public Health and Environment Riffle-run (CDPHE RR), and California’s Surface Water Ambient Monitoring Program (SWAMP). In **(B)** abbreviations are Large Wadeable (LW), Small Wadeable (SW), Boatable (B), Shocking (Sh), and Seine (Se).

### Accounting for Differences in Sampling Effort across Sampling Events and Improvements in Taxonomic Identifications through Time

Sampling effort and changes in the ability to identify taxa are remaining sources of variation and should be considered in the context of research questions because these factors influence sample abundances and probability of detection of any given organism. For instance, sampling effort varies across sampling events and is also correlated with time in some cases, which could lead to biased interpretation in certain analyses. For macroinvertebrates, these sampling effort variables include proportion sample identified (*PropID*), area sampled (*AreaSampled_m*), proportion of individuals in the lab identified to genus or species (*Gen_ID_Prop*), and the target number of individuals identified in a sample. *PropID* and *AreaSampledTotal_m* influence the number of individuals measured in a sample. As *PropID* and *AreaSampledTotal_m* increase, the total density of macroinvertebrates decreases, and as time proceeds *PropID* and *AreaSampledTotal_m* increase (Fig 8). Together, these trends suggest the potential that early years could be biased to large sample densities and later years could be biased to low sample densities, which would make it appear as though total densities decline over time.

**Figure 8.**
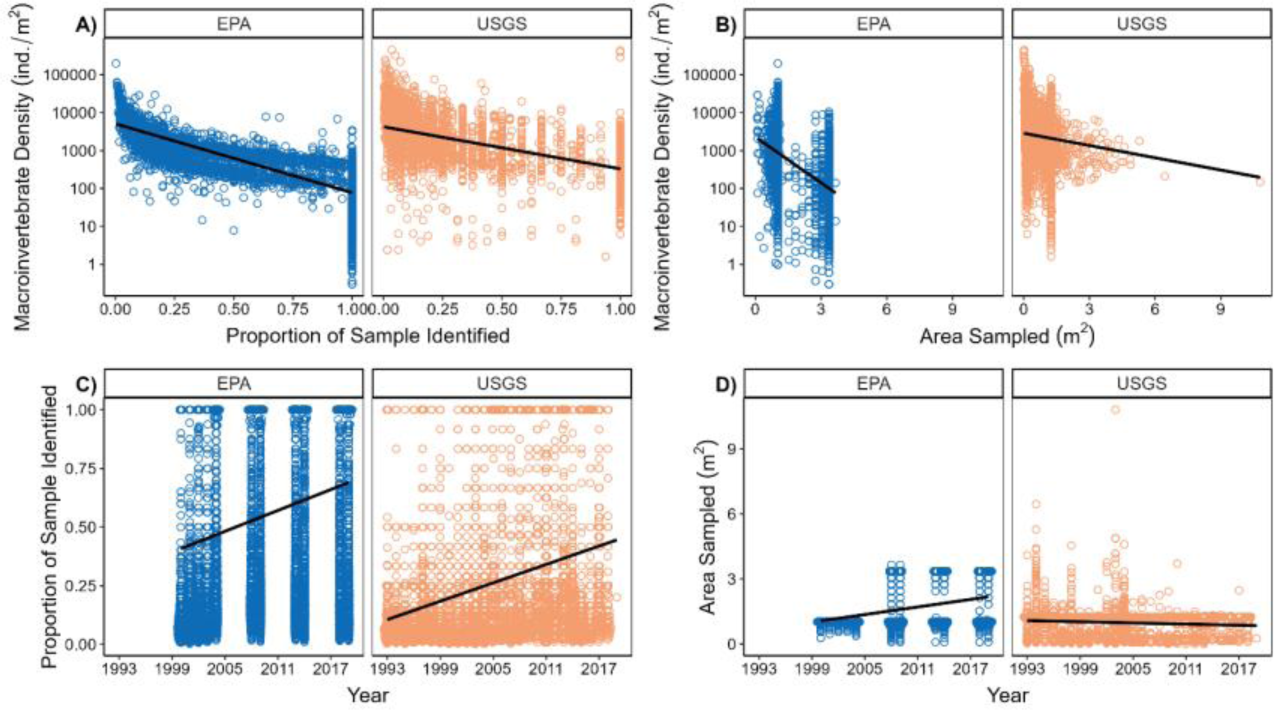
Associations among sampling effort variables, total macroinvertebrate densities, and time for both EPA and USGS datasets. Trend lines are simple linear relationships between sets of variables. Ignoring these variables could lead to spurious results in some instances.

The proportion of specimens (individuals) within samples identified to genus or species (*Gen_ID_Prop*) is a variable that describes the ability of taxonomists to identify macroinvertebrates. Within some orders, *Gen_ID_Prop* increases over time, likely indicating that taxonomists improved in their abilities to make identifications at the genus or species level, and as a result the probability of detection of these groups would appear to increase through time (Fig. 9). Users should take care to consider these patterns since they could lead to the false impression that new or more genera appear through time. Since most fish are identified to the species level, it is unlikely that the fish data have a similar issue.

**Figure 9.**
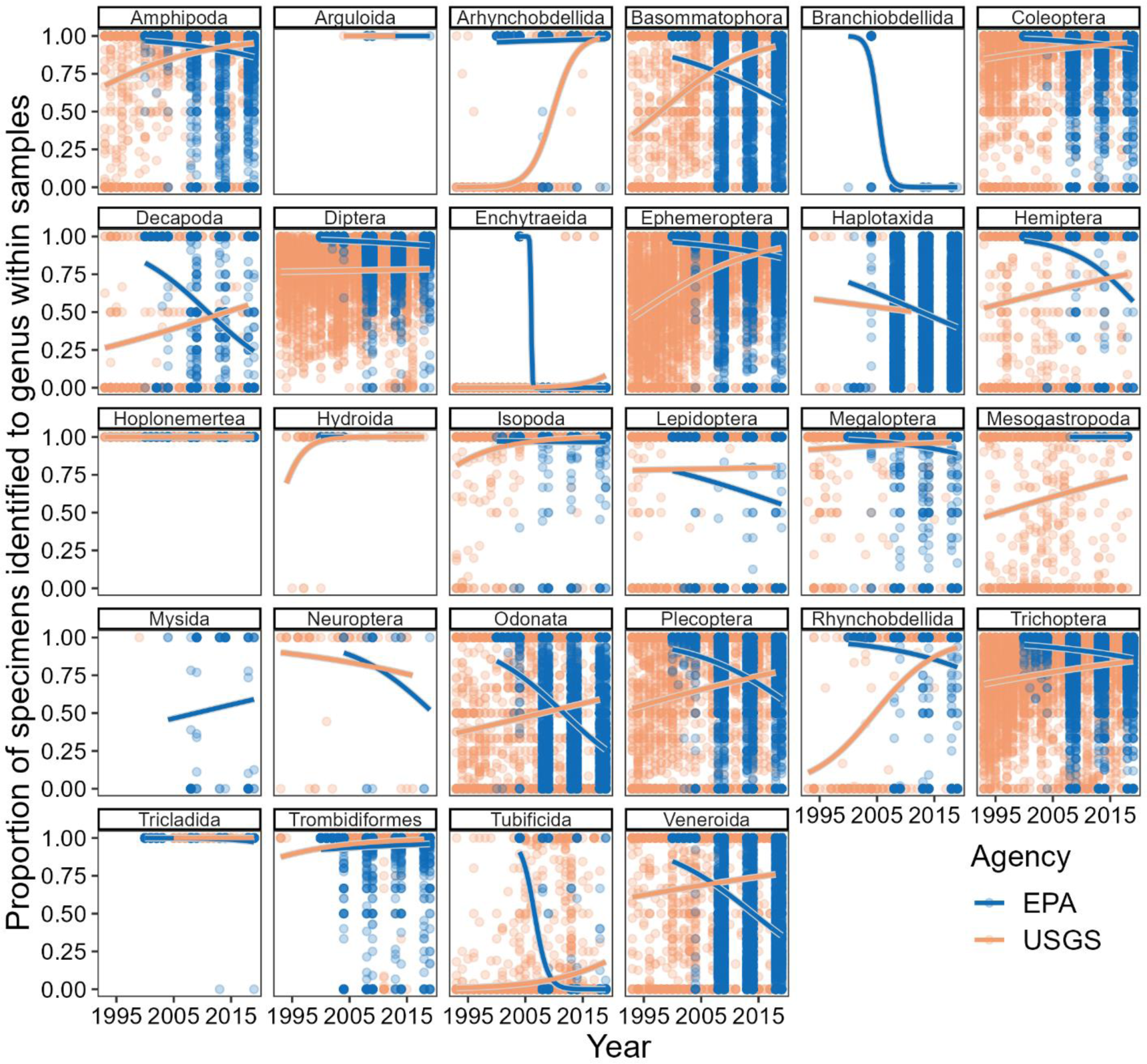
Changes in the ability of taxonomists to make genus or species identifications. Relationships between time and the proportion of specimens identified to genus within samples by order for EPA and USGS datasets. Positive trends may indicate that within a given order, taxonomists improved in their abilities to make identifications at the genus level.

Similarly, within some orders within the EPA dataset, proportion of specimens identified to genus decreases over time. One possible explanation for these trends could be the time spent identifying and enumerating WSA samples was greater than NRSA samples, and as a result the detection of these groups could appear to decrease through time.

The number of individuals within a sample is another component of sampling effort, and variation in this component is especially influential in the calculation of biodiversity metrics because sample richness increases with the number of individuals sampled. The target number of organisms identified within a sample varies across programs, agencies, time, and space, which introduces variation in output datasets. Target counts are typically 500 individuals for EPA samples, and they range from 100-600 for USGS samples. The ‘finsyncR’ package provides a parameter (rarefy = FALSE) within the `genInvertData()`to randomly resample samples to a specific individual count. When `rarefy = TRUE`, users set the organism count limit with the parameter `rarefyCount = 300` and samples containing at least that number are retained. Within the package, the rarefaction level defaults to 300 to balance the proportion of the samples dropped that failed to meet the rarefaction level with the addition of genera gained at the rarefaction level (Fig. 10). With a rarefaction level of 300, ∼91% of samples are retained, and with every 50 individuals identified, ∼1 genus is added to the average sample. So, lowering the threshold to 200 individuals would have removed ∼2 genera per sample and would have retained 94.5% of samples. Similarly, increasing the threshold to 400 individuals would have added ∼2 genera per sample but would have retained only 70.6% of samples.

**Figure 10.**
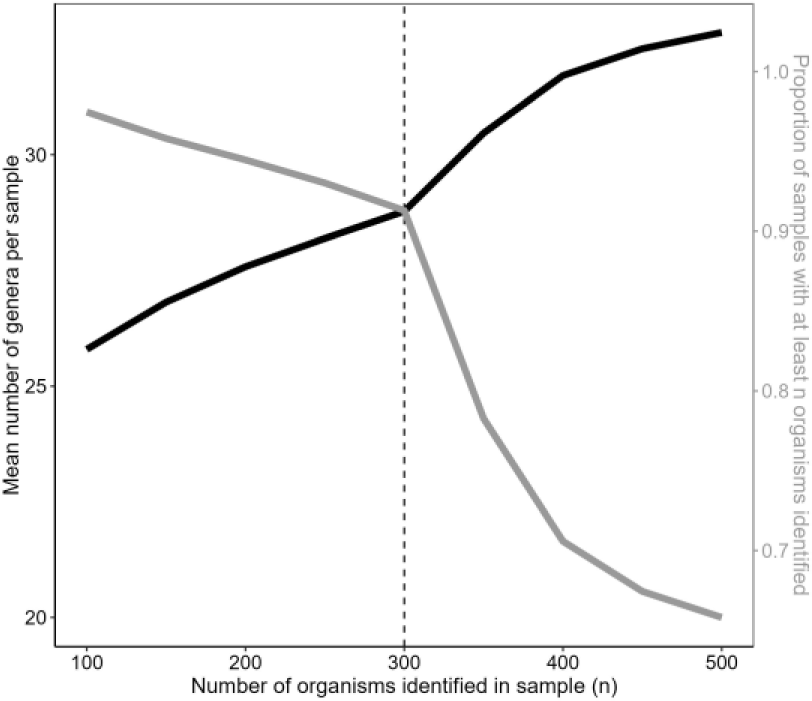
Justifying the use of 300 as the default rarefaction threshold. The relationship between mean number of genera gained, the proportion of samples with at least *n* number of organisms identified, and the number of organisms identified per sample.

Use `seed = 0` and provide an integer to consistently generate output from the function. This parameter is passed to set.seed() internally. The `rarefy = FALSE` parameter is not present in the `getFishData()` function because most methods do not incorporate a target count. An exception for fish is boatable streams in NRSA, which sampled until 20 stream widths or 500 organisms, whichever came first.

The fish sampling effort variables include reach length fished (*ReachLengthFished_m*), the sampling method employed (e.g. electrochocking, seining, snorkelling; *SampleMethod*), and standardized effort values (e.g. minutes of shocking, number of seine hauls, minutes snorkeling; *MethodEffort*). *MethodEffort* and *ReachLengthFished_m* influence the number of individuals measured in a sample. Here, we show a plot of their correlations with total number of fish within samples highlighting their potential importance (Fig. 11). However, unlike the macroinvertebrate sampling variables (Fig. 8), these fish sampling variables vary randomly through time. There could conceivably be instances in which fish sampling variables are important to consider in analyses given their wide variation and correlation with number of fish within samples.

**Figure 11.**
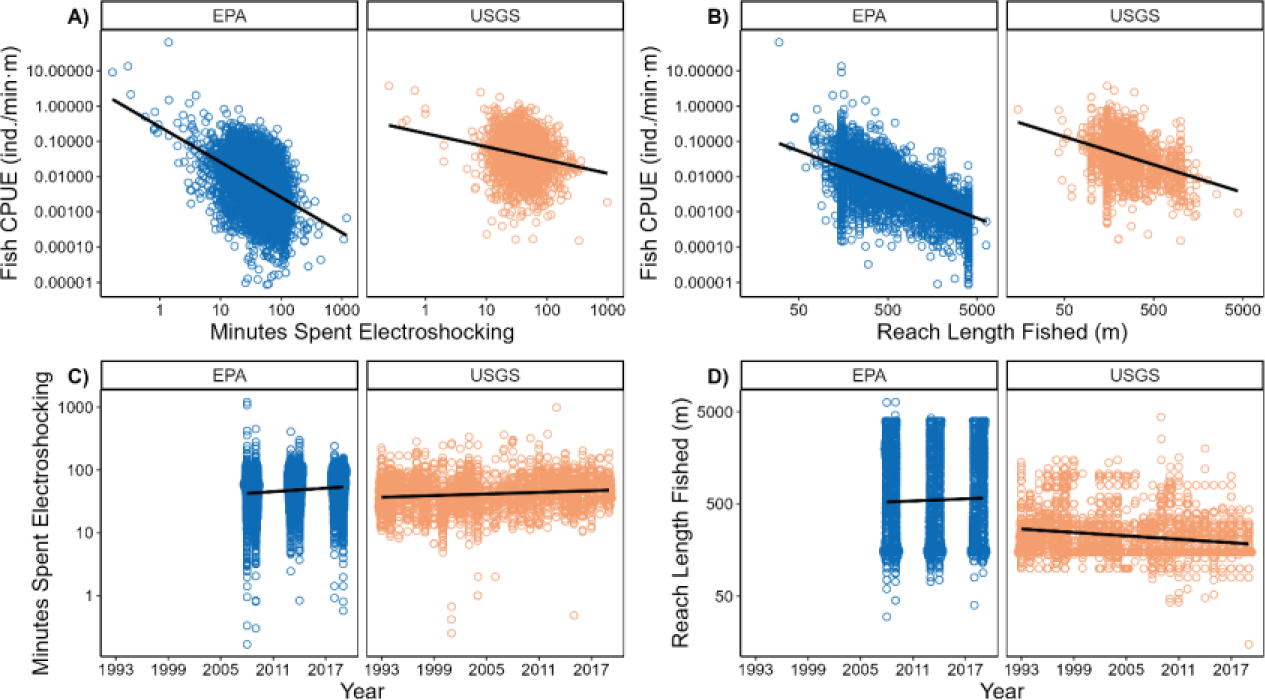
Associations among sampling variables, catch per unit effort (CPUE), and time for both EPA and USGS datasets. These associations suggest that users should consider these sampling effort variables. For instance, as *MethodEffort* and *ReachLengthFished_m* increase, the total number of fish decreases. However, unlike the macroinvertebrate sampling variables (Fig. 7), these fish sampling variables randomly vary through time.

## Usage Notes

### How to Use finsyncR

More information on how to implement code discussed in this manuscript is available in the Getting Started Vignette, which can be accessed with the following code `vignettes("Getting Started", "finsyncR")`. The metadata for the output datasets can be accessed within the package using the code `vignettes("Metadata", "finsyncR")`. We also provide a vignette on back calculating macroinvertebrate abundances and raw counts and rarefying raw counts which can be accessed using the code, `vignettes("BackCalculation", "finsyncR").`

### Disclaimers

The approaches provided in this package concerning the way the BioData and NRSA data are treated are not necessarily the same as the approaches used in other USGS and EPA publications or national assessments. So, replicating analyses using the processed data in this package may not produce identical results. For instance, multimetric indices produced from NRSA data from ‘finsyncR’ may not match multimetric indices in EPA national assessment reports.

The methods developed to join the EPA and USGS datasets described in this manuscript have not been designed with the intent of adding other fish and macroinvertebrate datasets. So, the appropriateness of these methods cannot be guaranteed for other datasets. The views expressed in this manuscript are those of the authors and do not necessarily represent the views or policies of the US Environmental Protection Agency.

## Code Availability

All code used within the ‘finsyncR’ package is publicly available. The ‘fincyncR’ package is stored via two different storage systems to provide users with different alternatives to accessing and using the data. The FigShare repository (DOI: 10.6084/m9.figshare.24938607) stores the fixed, original version of the package (version 1.0.0). The EPA GitHub repository (github.com/USEPA/finsyncR) stores the future updated versions of the R package. Users can install the finsyncR package directly from the GitHub repo using the following code: `devtools::install_github(“USEPA/finsyncR”, build_vignette = TRUE)`. Future updates may occur when new datasets are integrated.

Version 4.2.2 of R^32^ was used to build the ‘finsyncR’ package. Minimum versions of R and dependencies are listed in the ‘DESCRIPTION’ file (∼finsyncR/DESCRIPTION).

## Acknowledgements

We are indebted to the federal, state, and tribal biologists and contractors who planned these surveys, collected the samples in the field, and processed and identified samples. We thank them for their efforts in building these datasets that we consider a national treasure. We are especially grateful to Sarah Lehmann, Richard Mitchell, Amanda Nahlik, Dave Peck, and Alan Herlihy at the EPA and Daren Carlisle, Lindsey Boyle, Kelly Maloney, Taylor Woods, Travis Schmidt, Mike Meador, Scott Grotheer, Robert Zuellig, and Peter Ruhl at the USGS for the intellectual support of this work and the many conversations about the nuances of these data. Jason Rohr was a mentor for MBM, SLR, DKJ, EAB while portions of the current work were completed. This work was conducted as part of the Analyses of Contaminant Effects in Freshwater Systems: Synthesizing Abiotic and Biotic Stream Datasets for Long-Term Ecological Research Working Group supported by the John Wesley Powell Center for Analysis and Synthesis, funded by the U.S. Geological Survey.

## Author contributions

Michael B. Mahon: Conceptualization, Code development, Package maintenance, Methodology, Writing—original draft, Writing—reviewing and editing

Devin K. Jones: Conceptualization, Code development, Methodology, Writing—reviewing and editing

Ryan A. Hill: Code development, Writing—reviewing and editing Terry N. Brown: Code review, Writing—reviewing

Ethan A. Brown: Code development, Writing—reviewing and editing Stefan Kunz: Code development, Writing—reviewing and editing

Samantha L. Rumschlag: Conceptualization, Code development, Methodology, Project administration, Writing—original draft, Writing—reviewing and editing

## Supplemental Information

**Table S1.**
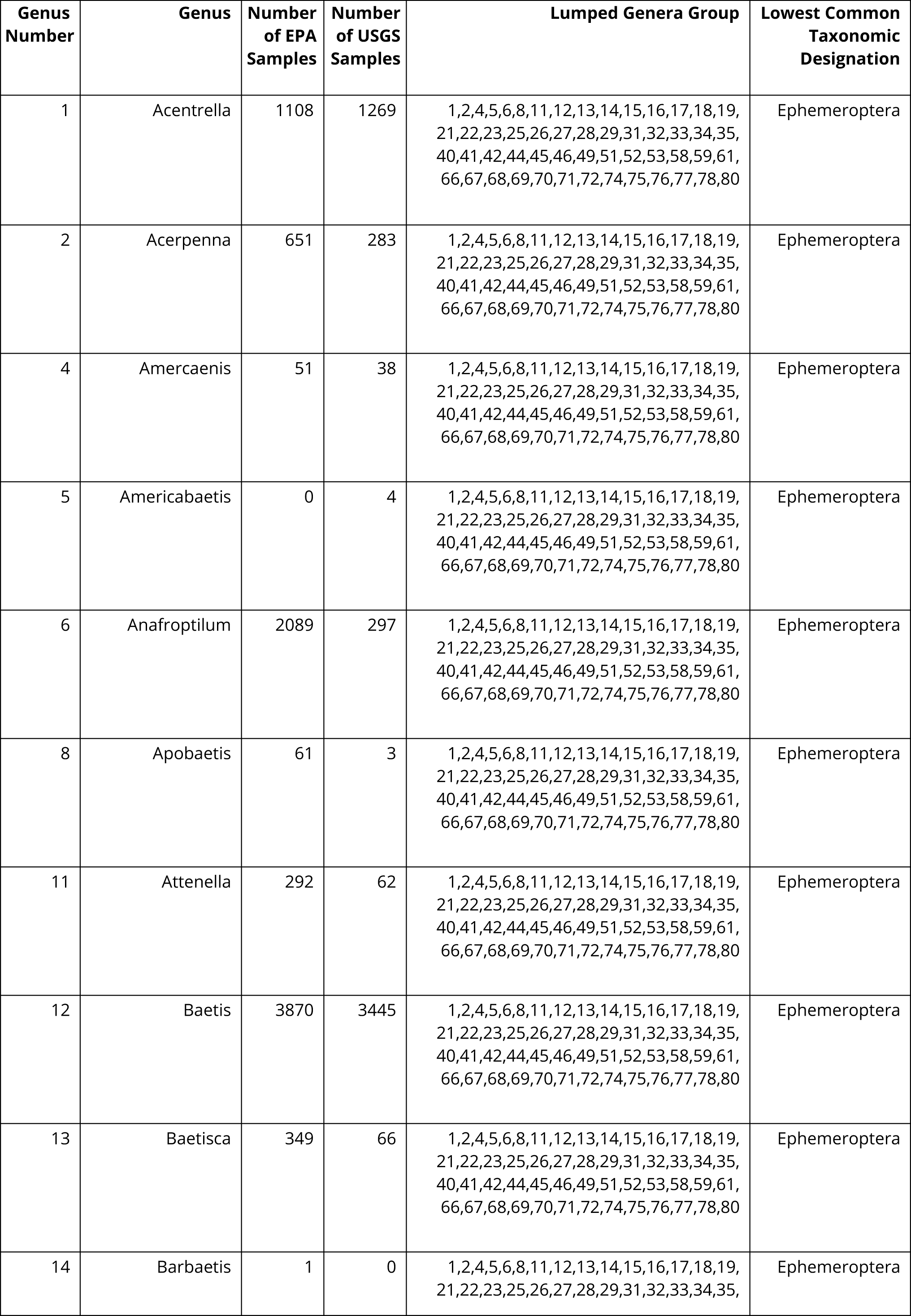
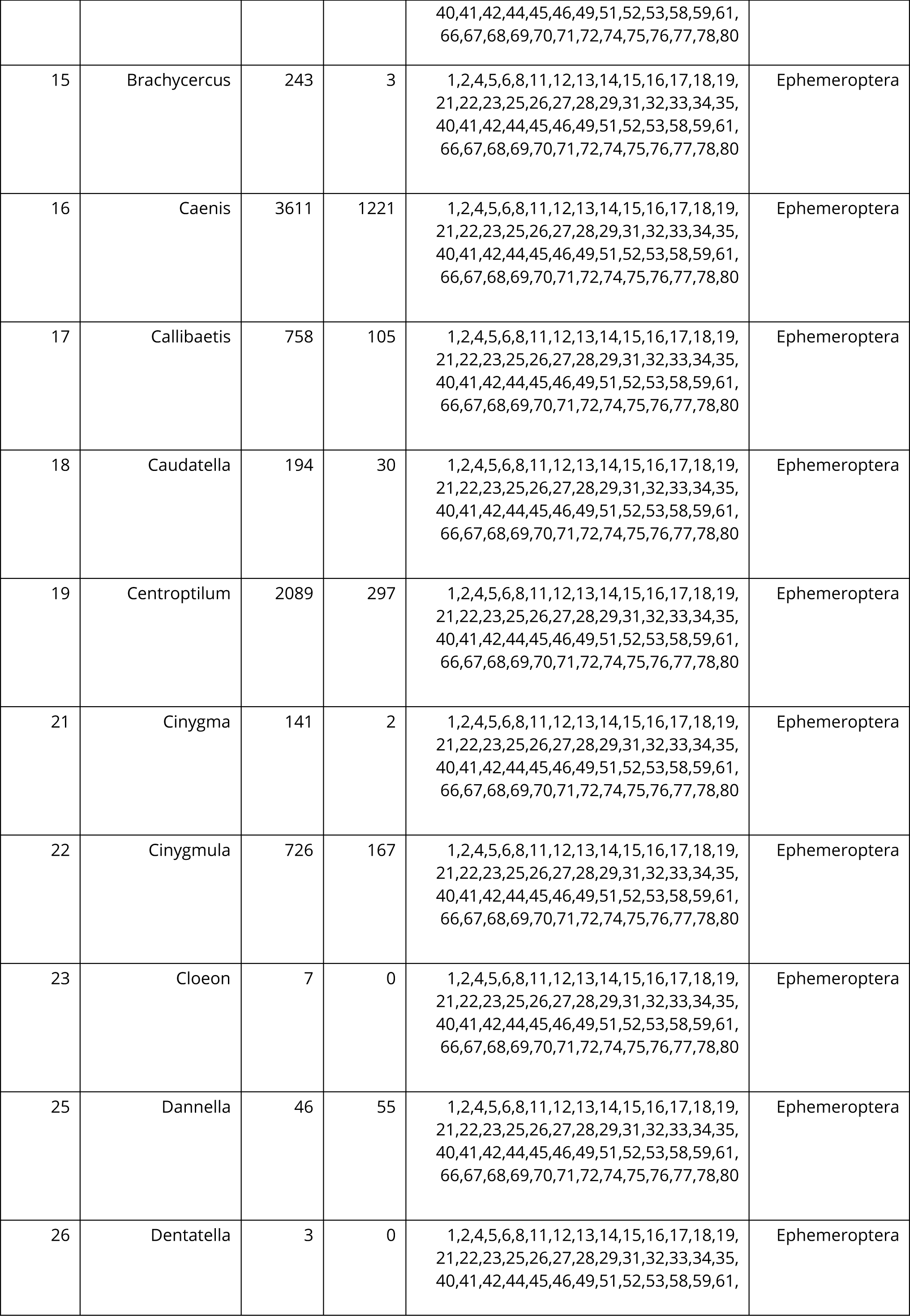
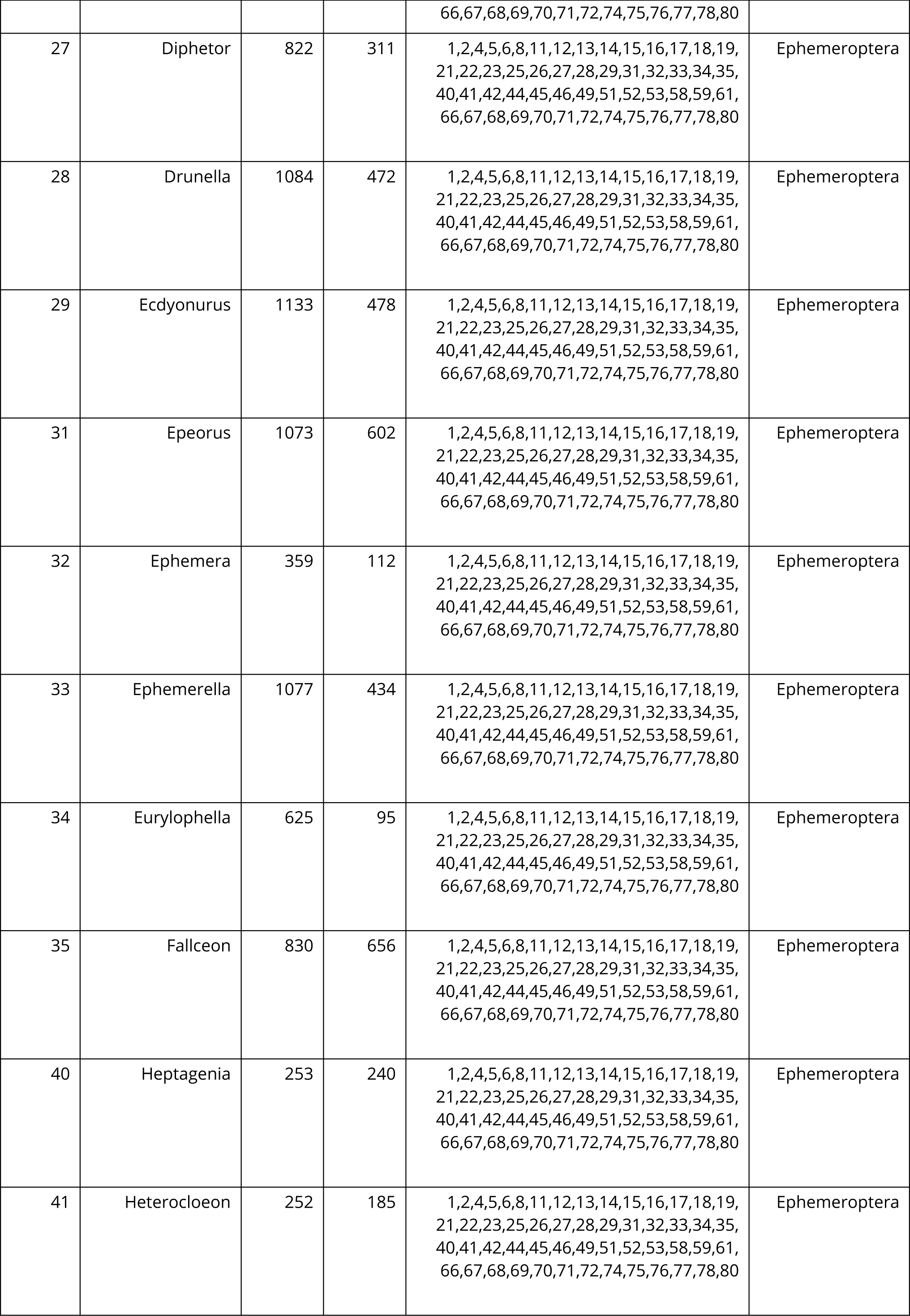
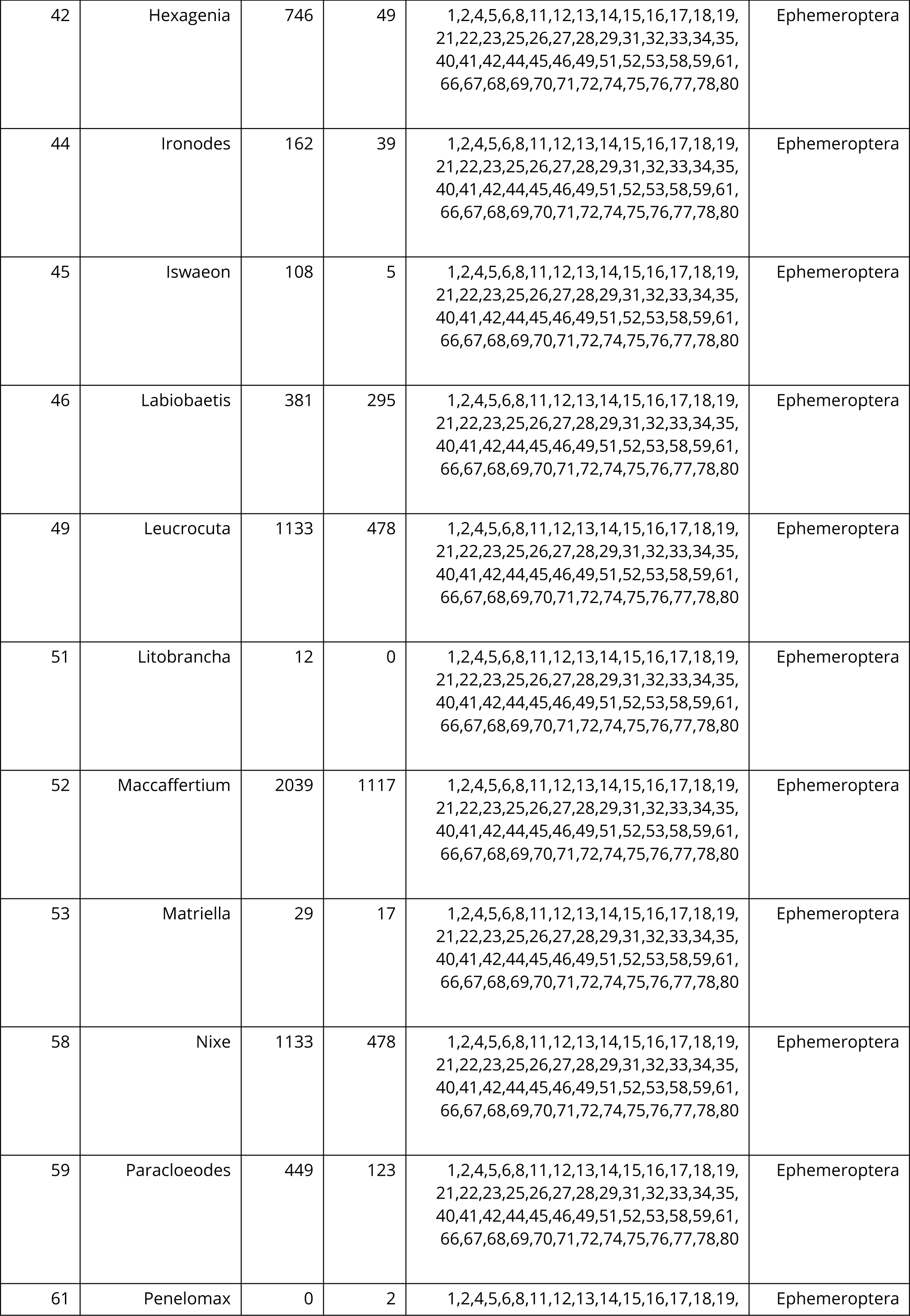
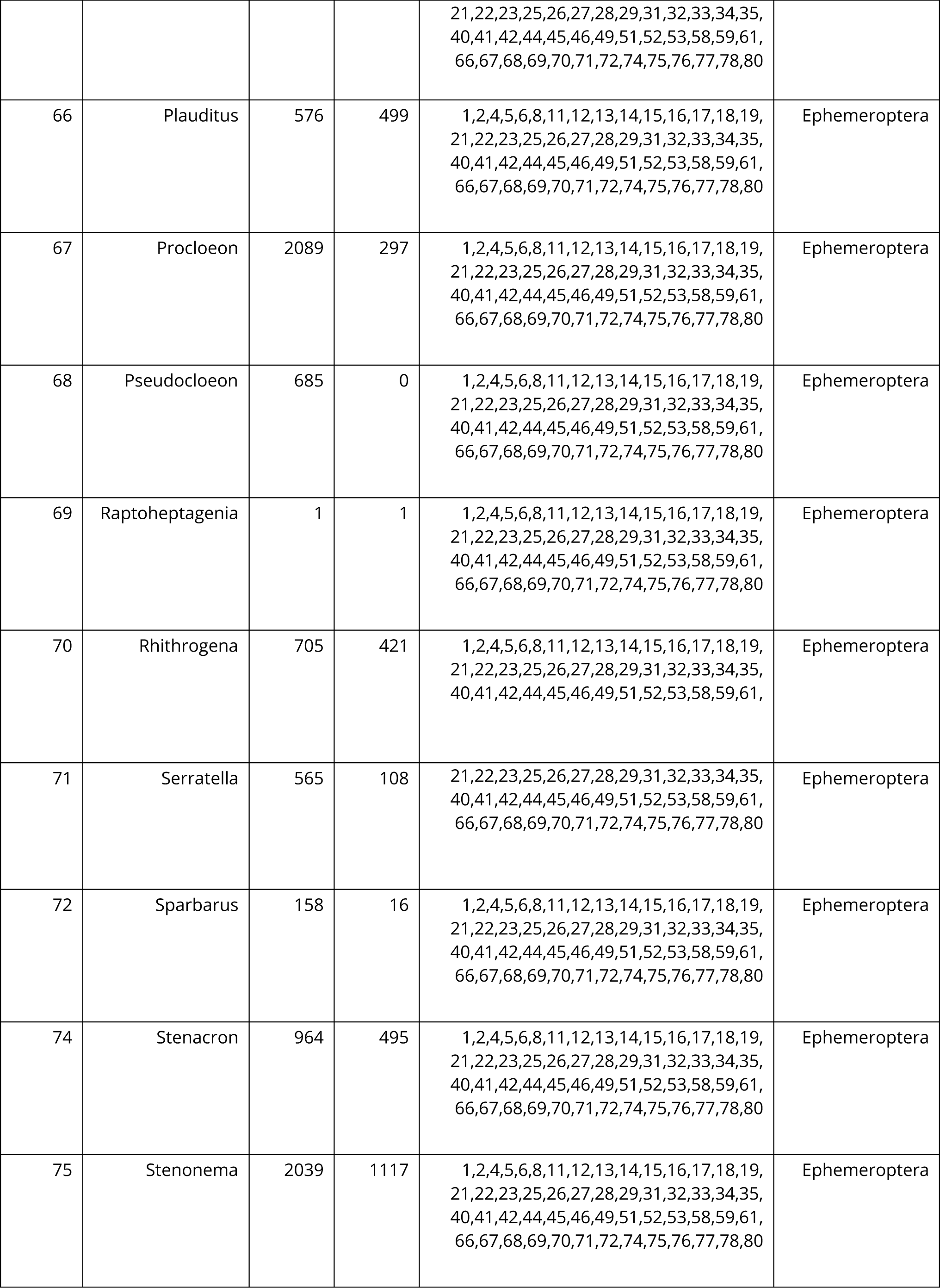
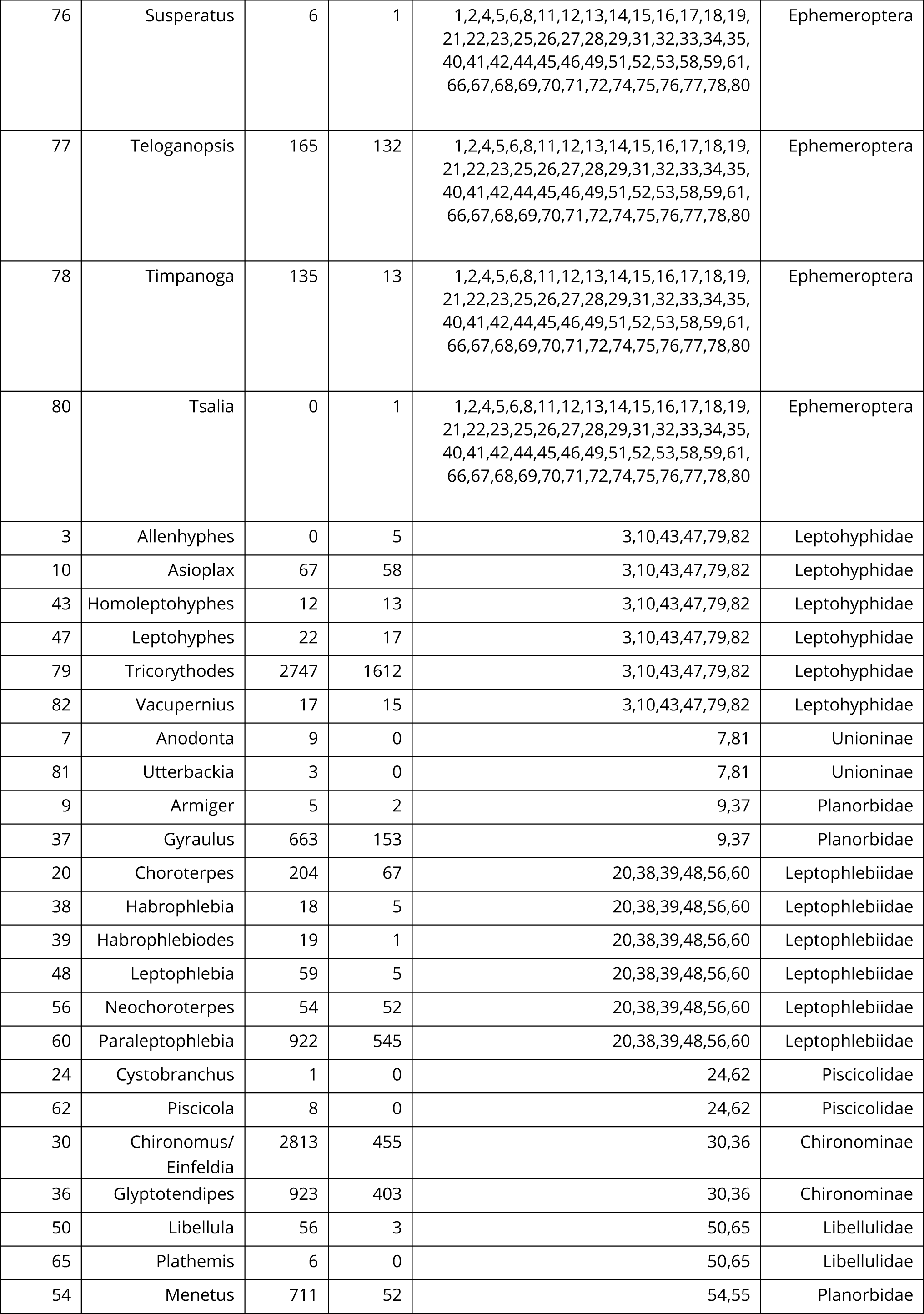
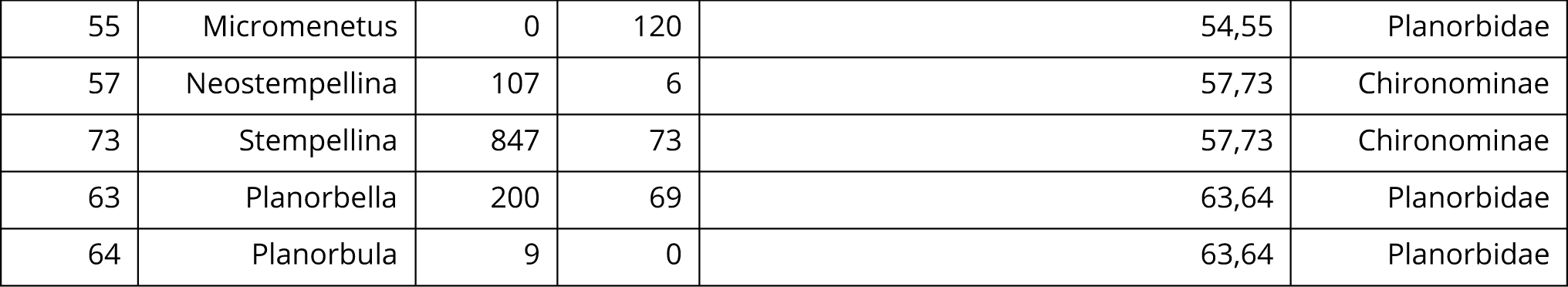
Table linking numbers from Figure 5 to genera names, associated lumped genera groups, and the number of samples in which each genus appears. Lumped genera groups account for changes in taxonomy through time. Some genera are found in very few samples, and users could consider dropping rare genera and recreating lumped groups depending on their data needs. The table also provides the lowest taxonomic designation common for the lumped group. This column is the taxonomic designation provided to the lumped group in the output from the `lumpRollUp()` function.

